# Mapping Haemagglutinin Residues Driving Antigenic Diversity in H5Nx Avian Influenza Viruses

**DOI:** 10.1101/2025.06.18.660338

**Authors:** Rebecca Daines, Jean-Remy Sadeyen, Pengxiang Chang, Munir Iqbal

## Abstract

Since its emergence in 1996, the H5 avian influenza virus (AIV) A/Goose/Guangdong/1/1996 (Gs/GD) haemagglutinin (HA) has evolved into over 30 genetically and antigenically distinct clades, including the widespread clade 2.3.4.4b. Vaccination is widely used in endemic regions to reduce poultry losses and zoonotic risk. However, the evolving antigenic diversity and global co-circulation of multiple clades challenges protective efficacy of poultry vaccines with poor antigenic matching to field strains, resulting in immune escape and vaccine failure. This study aimed to improve vaccine seed selection by identifying HA epitopes contributing to inter-clade antigenic differences.

Recombinant clade-representative viruses were generated using HA genes from circulating H5 AIVs via reverse genetics with A/Puerto Rico/8/1934 (PR8) internal and neuraminidase genes. Antigenic relationships were assessed using haemagglutination inhibition (HI) assays with homologous and heterologous chicken antisera. Antigenic cartography revealed a clear distinction of clade 2.3.4.4 from others and notable intra-clade diversity. Pairwise antigenic and genetic comparisons identified 48 putative antigenic residues. These were individually introduced into a candidate HA by site-directed mutagenesis, and antigenic influence assessed by HI using sera raised against the non-mutated HA.

Four residues R82K, A83T, T204I, and F229Y had significant antigenic effects, with three (R82K, T204I, F229Y) being novel. These findings demonstrate that combining serology and *in silico* residue analysis can identify key antigenic determinants.

This work highlights the need for precise antigenic matching in vaccine design and highlights the value of combining molecular and immunological tools to optimise vaccine seed selection against diverse and evolving H5 AIV strains.

**Importance:** The continued evolution of H5 avian influenza viruses (AIVs), particularly the Gs/GD lineage, poses major challenges for poultry disease control and zoonotic risk mitigation. Vaccine effectiveness is undermined by antigenic drift and the co-circulation of diverse clades, often leading to mismatches between vaccine and field strains. This study addresses the critical need to improve vaccine strain selection by identifying haemagglutinin (HA) residues driving antigenic variation across H5 clades. Using recombinant viruses, antigenic cartography, haemagglutination inhibition (HI) assays, and mutagenesis, 48 putative antigenic residues were identified, with four R82K, A83T, T204I, and F229Y having major antigenic effects, three of which were novel. These findings advance our understanding of H5 antigenic evolution and provide a framework for predicting vaccine performance. By integrating molecular and serological data, our work informs rational vaccine seed strain selection, contributing to more broadly protective vaccines and improved control of H5 AIV in poultry, while reducing the risk of zoonotic transmission.

## Introduction

Low pathogenic avian influenza (LPAI) viruses circulate naturally in wild aquatic birds, particularly in species from the orders *Anseriformes* and *Charadriiformes* [1]. These birds serve as reservoirs for all avian influenza virus (AIV) subtypes and typically show little to no clinical disease [1]. Their migratory behaviour drives global virus distribution, with occasional spillover of LPAI subtypes into domestic poultry populations. In poultry, these viruses can establish persistent circulation and cause mild to severe disease [2]. Notably, highly pathogenic avian influenza (HPAI) viruses can emerge when LPAI H5 or H7 subtypes infect galliform poultry, acquiring mutations in the haemagglutinin (HA) that enable systemic infection, leading to increased severity and mortality [3]. These HPAI strains can spill back into wild bird populations, often causing subclinical or less severe disease, and may be reintroduced into naïve poultry flocks [4, 5].

A novel outbreak of H5 HPAI with high morbidity was first detected in domestic geese in China in 1996, resulting in the emergence of the H5 HA A/Goose/Guangdong/1/1996 (Gs/GD) lineage [1]. This strain differed from earlier, sporadic H5N1 HPAI viruses [6, 7]. Since its emergence, the Gs/GD lineage has diversified into 10 genetically distinct clades (0 – 9), each with multiple subclades [8]. In 2013, clade 2.3.4.4 emerged and has since become dominant in H5 outbreaks, particularly in Europe, Asia, and the Middle East, often co-circulating with reassorted neuraminidase (NA) subtypes, collectively termed H5Nx. This clade further diversified into subclades 2.3.4.4a – h, initially named based on their geographical origins [9]. September 2022 to February 2023 saw unprecedented international loss of domestic and wild bird populations from HPAI H5N1 of clade 2.3.4.4b, presenting as multiple reassortments, H5N1, N2, N4, N5, N6 and N8 [10]. Most notably, the host range expanded into domestic and predatory mammals, aquatic porpoises such as whales or dolphins and co-habiting aquatic mammals including seals, significantly increasing the risk of zoonotic potential.

Vaccination remains a key strategy to reduce both the socio-economic impacts AIV infections and the zoonotic risk in endemic regions. However, the rapid genetic and antigenic evolution of AIVs, particularly, H5Nx viruses from clade 2.3.4.4, complicates effective vaccine seed strain selection. Poor antigenic matching may result in suboptimal protective immunity or drive immune escape, accelerating viral evolution [11]. Vaccine strain updates are typically based on haemagglutination inhibition (HI) assays that measure the cross-reactivity of vaccine-induced sera against circulating viruses [12]. When cross-reactivity drops below a defined threshold, a new seed strain is selected to restore vaccine efficacy and reduce the risk of vaccine failure. While whole inactivated virus vaccines are favoured for their low cost and scalability, they may quickly become antigenically outdated and fail to cover the breadth of circulating diversity. In response, in silico approaches, including synthetic antigen design, chimeric constructs, and multivalent formulations have been developed to expand vaccine coverage but are often constrained by production complexity and time [13].

The antigenic diversity and co-circulation of H5Nx clades often confounds efficient vaccination regimes or challenges accurate antigenic matching. Therefore, this research sought to adopt *in silico* methods to identify the residues driving the antigenic disparity between the clades of H5Nx. The identification of these residues can not only aid our understanding of the antigenic variability of H5Nx AIVs but direct more accurate and regionally targeted vaccine seed selection and alleviate the socio-economic burden faced by the poultry industry.

## Results

### Genetic variability within and between H5Nx clade 2.3.4.4 subclades is significantly greater than in other clades

Representative candidate viruses were selected from both current and recently active clades of the Gs/GD lineage of H5Nx AIVs, based on WHO/FAO/OIE biannual reports on antigenic characterisation and candidate vaccine virus (CVV) preparedness [8, 9]. Virus selection was guided by these reports as well as the revised nomenclature established in 2015 [14]. However, due to the rapid and ongoing antigenic diversification of clade 2.3.4.4, nomenclature updates have become outdated. Therefore, representatives from this clade were additionally selected based on WHO and CDC risk assessment reports and current CVVs [10, 15].

A total of 22 viruses were selected, spanning clades 1.1.1 to 2.3.4.4h. Full-length haemagglutinin (HA) sequences were retrieved from public databases, including NCBI [16] and GISAID [17] (**Table 1**). Sequences were trimmed to the mature open reading frame, aligned, and quality checked. For clarity, strain names are hereafter referred to by shortened designations.

**Table 1:**
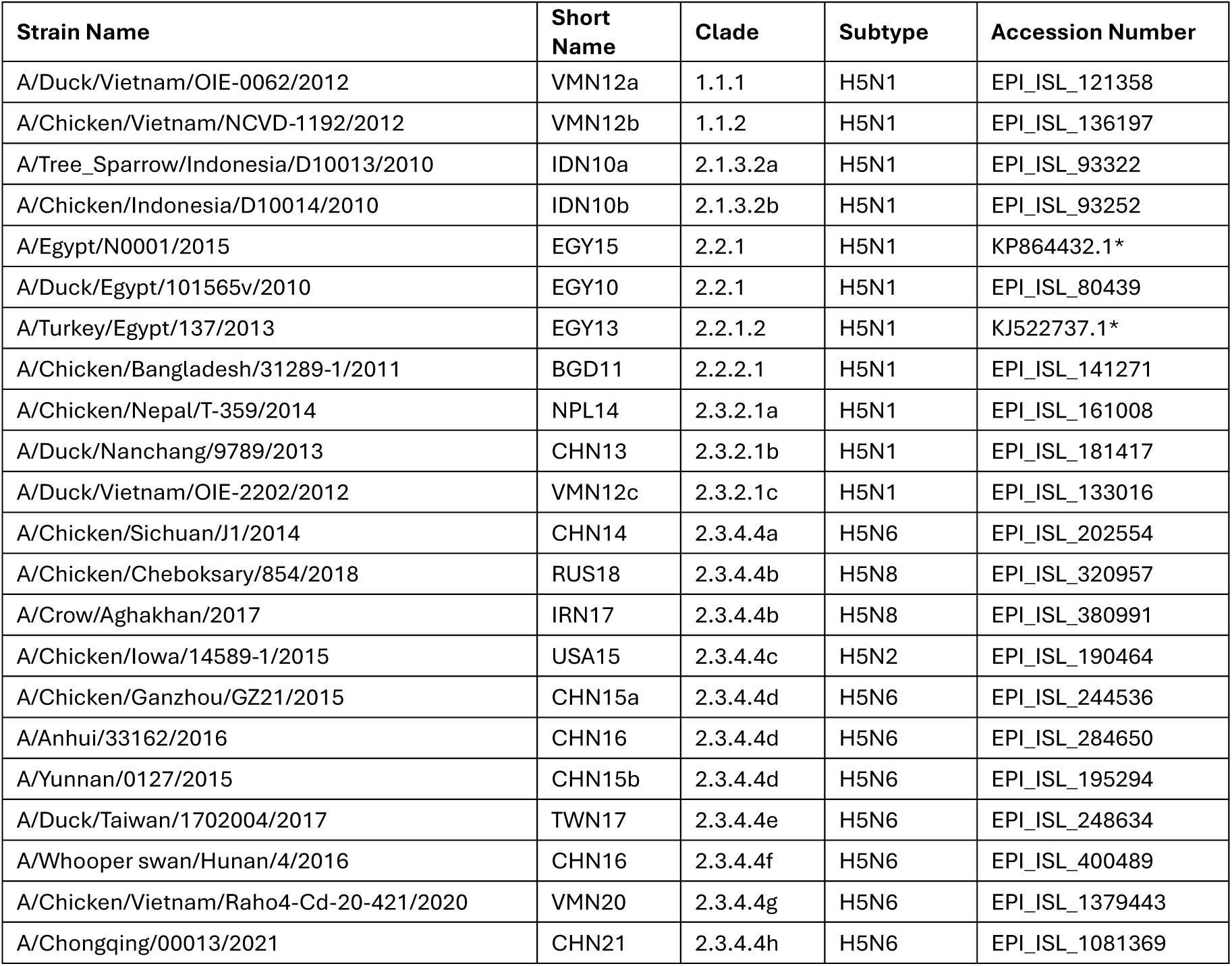
Representative H5Nx strains with associated clade and subtype chosen for this study. Short names adopt ISO country three-letter codes with the year of isolation. Strains sourced from GISAID except for (*) sourced from NCBI.

To confirm clade allocation and assess genetic variability while excluding the influence of synonymous codon usage, the selected HA sequences were translated into amino acid sequences and analysed phylogenetically. These were compared against WHO/OIE/FAO- assigned reference strains according to the established nomenclature system [10, 14]. A maximum-likelihood phylogenetic tree was constructed using the GTR+G substitution model, selected based on the lowest Bayesian Information Criterion (BIC) among best-fit models.

The tree was rooted to the clade 0 Gs/GD strain, with the assumption of equal evolutionary rates across lineages **(Figure 1**). Conserved amino acid residues are annotated on the branches where they are shared across all sequences within the respective clade or subclade.

**Figure 1:**
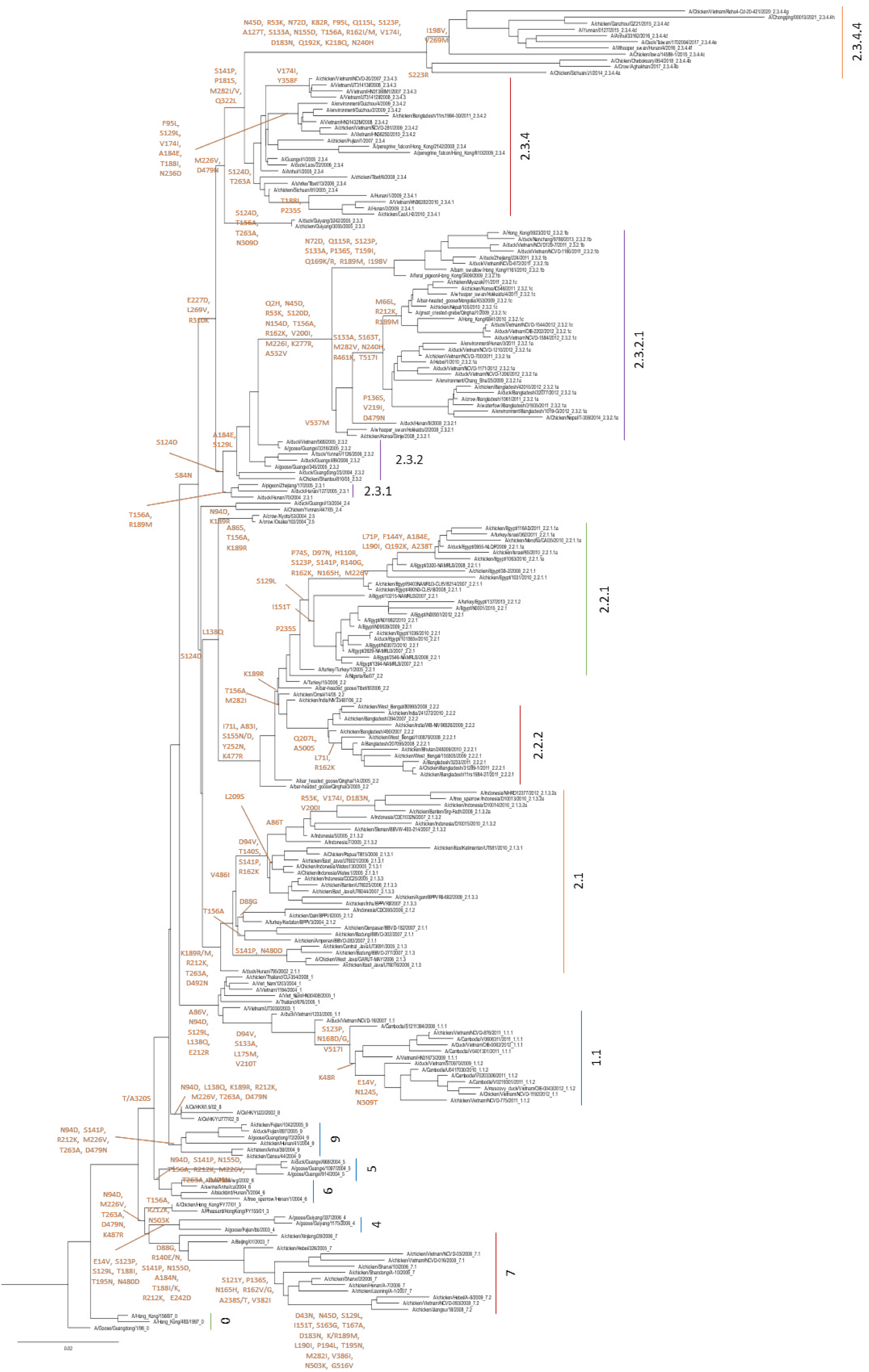
Maximum likelihood phylogenetic tree of Gs/Gd lineage H5Nx HA amino acid sequences. Sequences sourced from WHO/OIE/FAO nomenclature (clade 1 - 9, without 2.3.4.4) and selected clade 2.3.4.4 strains from GISAID and the 2020 CDC risk assessment for clade 2.3.4. with 4 viruses [1]. Conserved amino acid mutations shown on branch, all preceding strains containing the mutation. Tree rooted to A/Goose/Guangdong/1/1996. Tree constructed using IQ-Tree [2, 3] using JTT+G model selected for best fit with a fixed rate of heterogeneity across sites and 90% deletion criteria [4]. Mature H5 numbering used.

Clade 2.3.4.4 exhibited the highest level of intra-clade genetic variation, featuring 17 unique conserved residues across its subclades. This was followed by the now-extinct clade 7.2 with 15 conserved residues, and the geographically restricted clade 2.3.2.1 with 11. Notably, significant genetic diversification was also observed within clade 2.3.2.1 itself; subclade 2.3.2.1b formed a distinct branch with an additional nine conserved residues, while subclades 2.3.2.1a and 2.3.2.1c shared six further conserved sites, with 2.3.2.1c exhibiting an additional three unique residues.

To further investigate the extent of genetic diversity among the selected strains and support downstream analyses, pairwise amino acid differences across the mature HA sequences (excluding the signal peptide) were calculated **(Figure 2)**. A heatmap was generated to visualise these variations, with colour gradients ranging from purple (low divergence) to yellow (high divergence), representing the degree of pairwise sequence dissimilarity.

**Figure 2:**
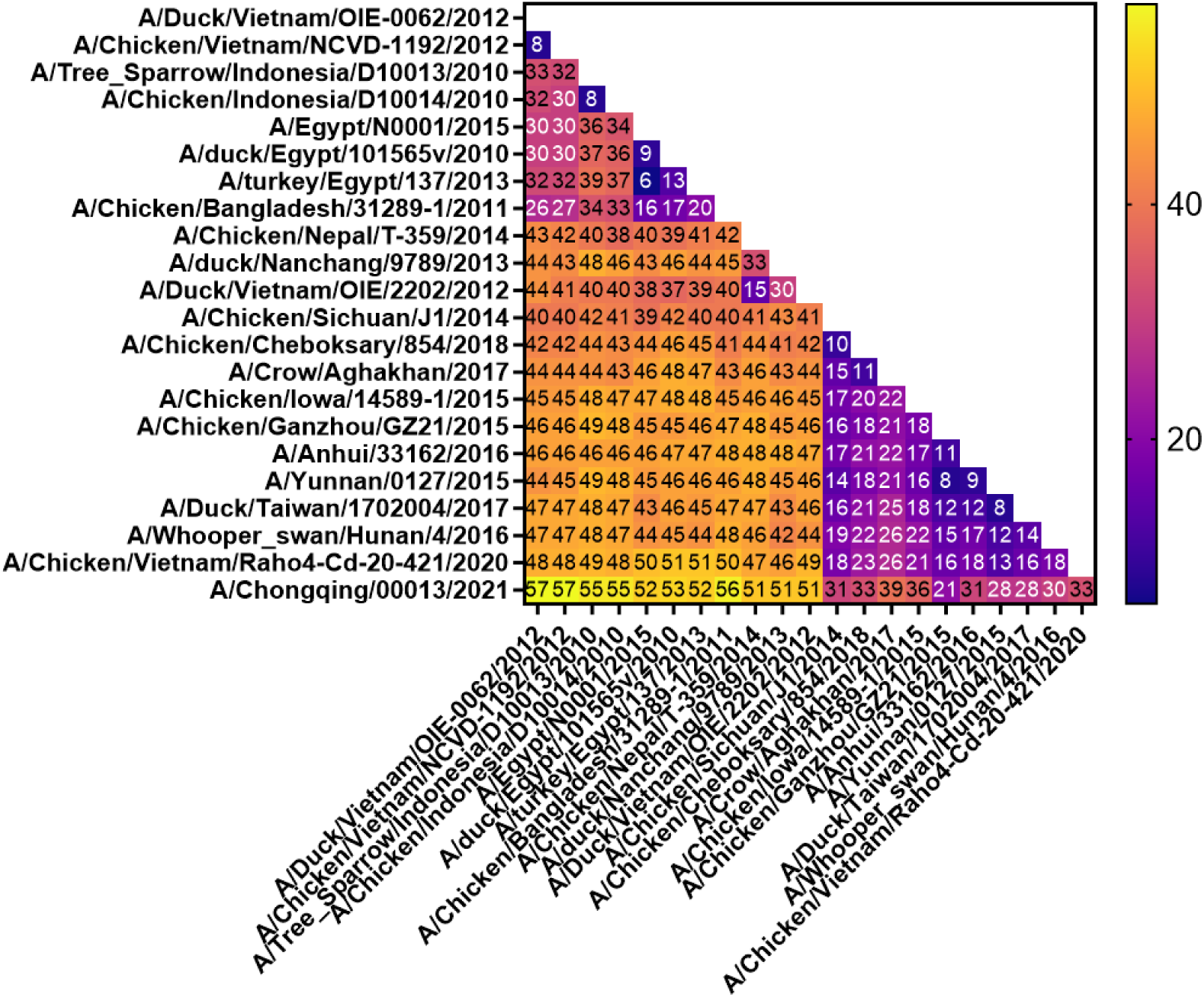
Heatmap of pairwise amino acid variances of the mature haemagglutinin (HA) sequence between strains. Pairwise differences were compiled in MEGA X and presented as a heatmap using GraphPad [5].

Amino acid variation among the selected strains ranged from six to 57 differences. Lower pairwise variation was generally observed between viruses from closely related clades and similar geographic origins; six differences were observed between EGY13 and EGY15 (clades 2.2.1.2 and 2.2.1, respectively), and eight differences between VMN12a and VMN12b (1.1.1 and 1.1.2) as well as IDN10a and IDN10b (2.1.3.2a and 2.1.3.2b).

In contrast, the highest pairwise divergence (57 differences) was observed between CHN21 (clade 2.3.4.4h) and VMN12a (clade 1.1.1), reflecting the substantial genetic distance between these clades. Within clade 2.3.4.4 itself, intraclade variation ranged widely from eight to 39 differences. The lowest variation was between CHN15a and CHN15b (both 2.3.4.4d), while comparisons between CHN15b (2.3.4.4d) and TWN17 (2.3.4.4e) also showed limited divergence. The greatest variation within clade 2.3.4.4 was consistently associated with CHN21 (2.3.4.4h), which differed by at least 21 amino acids from all other 2.3.4.4 representatives.

Antigenic diversity in AIVs is predominantly driven by amino acid changes within the HA1 region of the HA protein, located upstream of the proteolytic cleavage site. This region encompasses the receptor-binding domain (RBD), a critical target for neutralising antibodies. Due to its immunological importance, the HA1 region was extracted from each HA sequence for focused analysis of genetic variation and used in subsequent antigenic mapping **(Table 2)**.

**Table 2:**
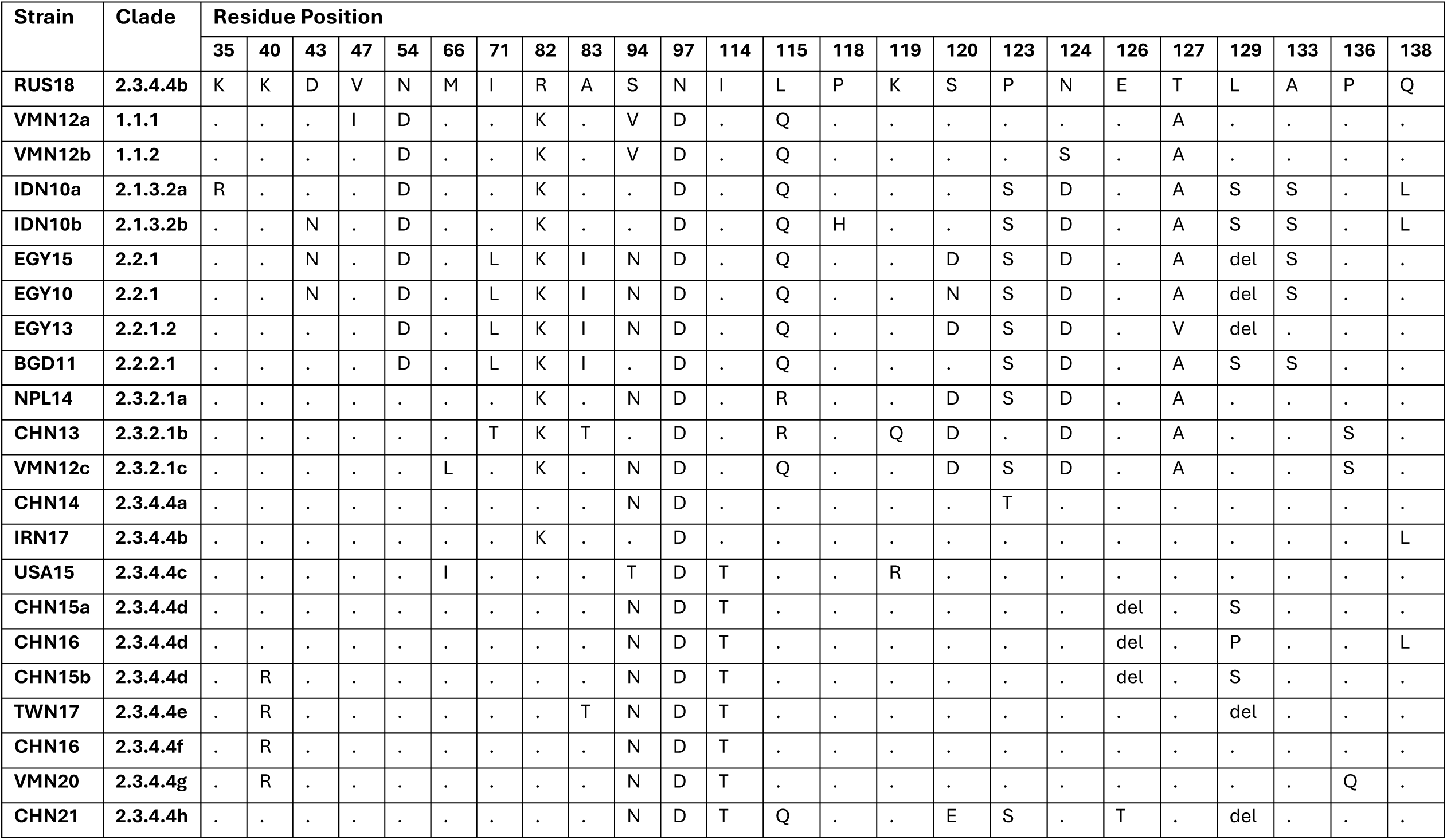
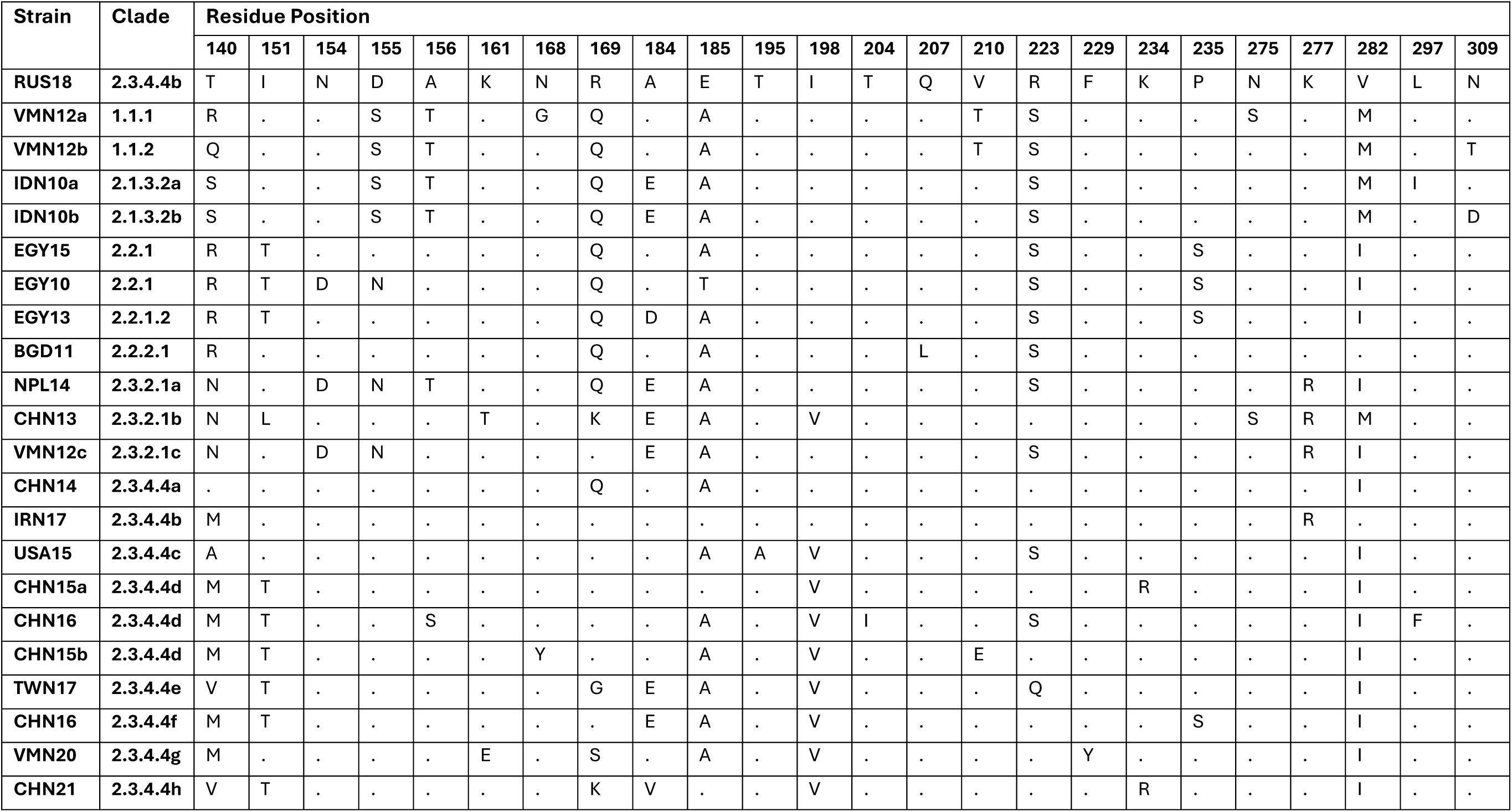
Amino acid variance of HA1 between the representative clade strains. Residue positions with variances between the strains are shown in comparison to the Rus18 2.3.4.4b strain. Deletions indicated with (del). Mature H5 numbering used [6, 7]. Amino acids are stated in their single-letter code.

A total of 48 amino acid variations were identified within the mature HA1 region (excluding the signal peptide and cleavage site) of the selected strains. Of these, 21 positions (43.75%) exhibited two amino acid variants, another 21 positions (43.75%) showed three variants, and the remaining six positions (12.5%) had four or more variants when compared to the RUS18 reference sequence. The most variable sites, residues 94, 120, 129, 140, 169, and 184, exhibited four, four, four, eight, five, and four amino acid variations, respectively.

Glycosylation of HA can also influence antigenic diversity. Therefore, sequences were analysed for potential N-linked glycosylation using the NetNGlyc server [18] (**Figure 3**). Glycan sites were predicted based on the presence of the canonical sequon Asn-X-Ser/Thr (where X is any amino acid except proline), with a confidence threshold of 0.5 (mean score from nine neural networks). No O-linked or C-linked glycosylation sites were identified.

**Figure 3:**
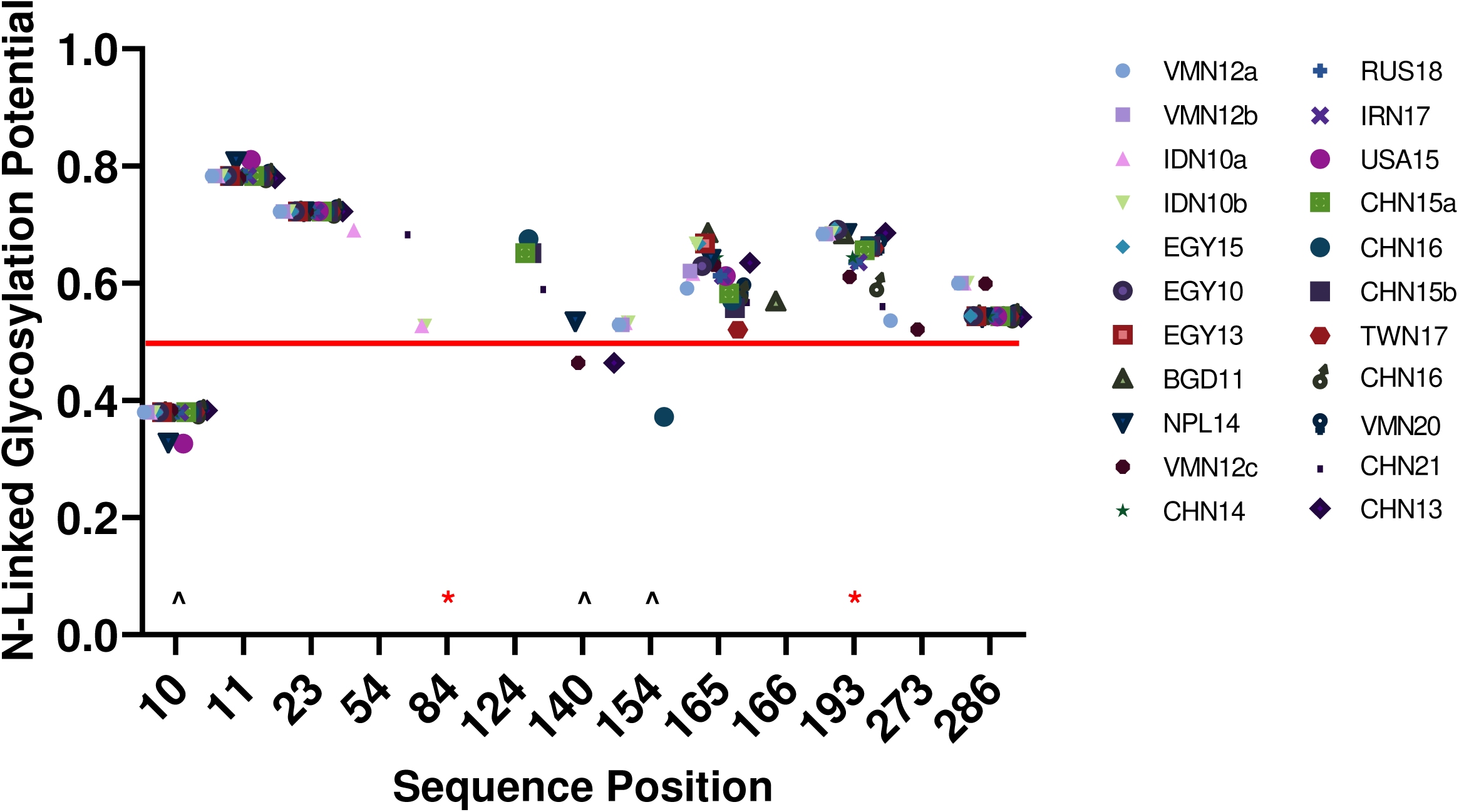
N-linked glycosylation prediction of study haemagglutinin (HA) sequences. The ‘potential’ score is the mean output of nine neural networks. Any symbol crossing the default threshold of 0.5 (red line) represents a predicted glycosylated site (if it occurs in the required sequon Asn-X-Ser/Thr without proline at X). Sequons with Pro present but achieved a positive confidence are indicated with (**\***). Sequons without Pro but achieve a negative confidence are indicated with (**^**). Mautre H5 numbering used. Prediction run using NetNGlyc [8].

Two predicted N-linked glycosylation sites, at positions 165 and 286, were conserved across all sequences. Although position 193 contains the required Asn-X-Ser/Thr sequon, the presence of proline at the X position in nearly all sequences disrupts glycosylation, with the exception of EGY13 and USA15. A similar disruption is observed at position 84 in IDN10a and IDN10b due to a proline residue within the sequon.

Additional sites showed positive glycosylation predictions at position 54 (IDN10a and CHN21), 124 (CHN15a, CHN15b, CHN16), 140 (NPL14), 154 (VMN12a, VMN12b, IDN10a, IDN10b), 166 (BGD11), and 273 (VMN12a and CHN13). While positions 140 (CHN13, VMN12c) and 154 (CHN16) contain intact sequons, they did not exceed the confidence threshold of 0.5, suggesting a lower likelihood of glycosylation.

### All subclades within clade 2.3.4.4 are significantly antigenically distinct from other co-circulating or recently circulating clades and exhibit considerable intra-clade diversity

To identify correlates of antigenicity among clades, the selected strains were assessed for cross-reactive neutralising antibody responses by measuring their ability to inhibit the binding of heterologous strains. Recombinant viruses were generated using reverse genetics (RG), incorporating the HA gene from each selected strain and the neuraminidase (NA) and internal genes of A/Puerto Rico/8/1934 (PR8). HA sequences were modified to remove the polybasic cleavage site and adapt non-coding regions for compatibility with the pHW2000 eight-plasmid bidirectional expression system. Polyclonal antisera were raised in chickens vaccinated with inactivated, concentrated, and adjuvanted recombinant viruses. Per strain, four to ten birds were inoculated at one day of age and boosted at seven days post-vaccination. Serum samples were collected every seven days, and antibody titres were assessed by HI assay. The final bleed at 42 days post-vaccination was used for cross-strain antigenic comparisons **(Table 3)**. HI assays were additionally validated by virus neutralisation tests (VNT), with a sample of HI titres compared to VNT titres **(Figure S1)**.

**Table 3:**
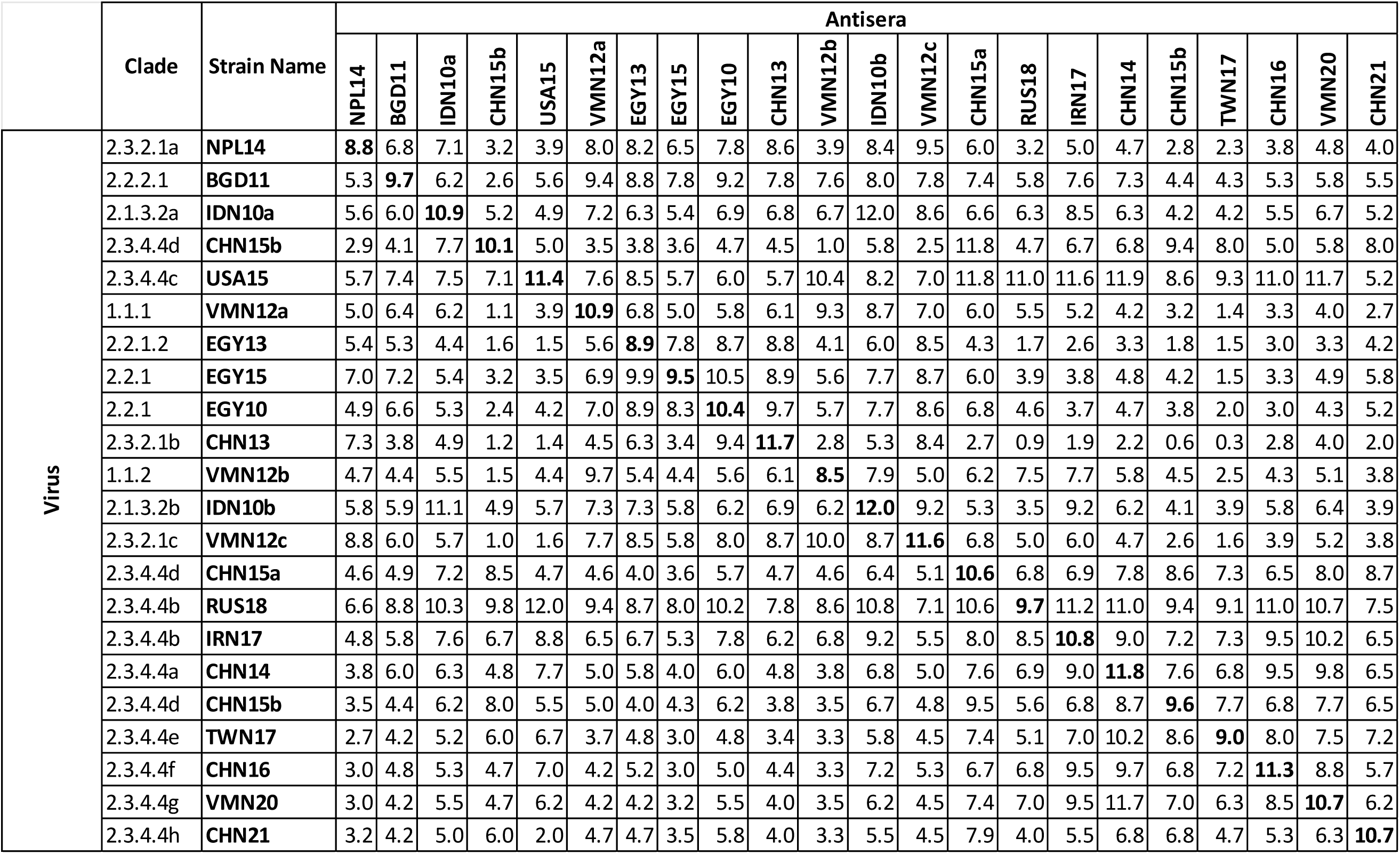
Geometric mean titres (GMT) of haemagglutination inhibition assays (log_2_) using homologous (**bold**) and heterologous whole virus antisera raised in chickens. Columns and rows represent sera and viral antigens, respectively.

Geometric mean titres (GMTs) were calculated using data from four to ten biological replicates (individual bird sera) and three technical replicates per strain. A titre of ≥5 log₂ (equivalent to 32 haemagglutinating units (HAU)) is considered protective [19–21]. All homologous titres (i.e., where the test antigen matched the immunising strain) exceeded this threshold substantially, ranging from 8.5 to 12 log₂ (384 to 4096) HAU. As anticipated, heterologous titres varied widely. Several were below the protective threshold, highlighting limited cross-protection between certain clades. Interestingly, RUS18 (clade 2.3.4.4b) and USA15 (clade 2.3.4.4c) were the most broadly reactive strains, with their HA inhibited by all heterologous sera tested, showing titres up to 11.2 and 11.9 log₂ (2432 and 3896 HAU), respectively. In contrast, CHN13 was the most antigenically distinct virus, showing cross-reactive inhibition with sera from only four heterologous viruses: NPL14, EGY13, EGY10, and IDN10a.

Although most reciprocal HI titres between antigen/antiserum pairs correlated well, some showed considerable discrepancies. For instance, sera raised against RUS18 tested against the CHN15a virus yielded titres of 6.8 log₂ (109 HAU), whereas CHN15a sera against RUS18 produced a markedly higher titre of 10.8 log₂ (1728 HAU). A total of 48 antigen/antiserum pairs exhibited differences greater than 3 log₂ , with the majority associated with RUS18 (17 pairs), USA15 (11 pairs), and CHN13 (9 pairs).

To visualize antigenic cross-reactivity among clades, an antigenic map was generated using the GMTs from Table 3. Antigens and sera were plotted in both two-dimensional (2D) and three-dimensional (3D) spaces, where spatial proximity reflected antigenic similarity, closer points indicated greater cross-reactivity. Initial mapping and optimisation were performed using ACMACS [22], followed by refinement and plotting in RStudio with the Racmacs package. An optimal minimum column basis (OMCB) of 128 and 500 optimisation runs were applied to minimise error and stress. Additional relaxation and randomization through dimensional annealing confirmed the robustness of the final maps (**Figure 4)**.

**Figure 4:**
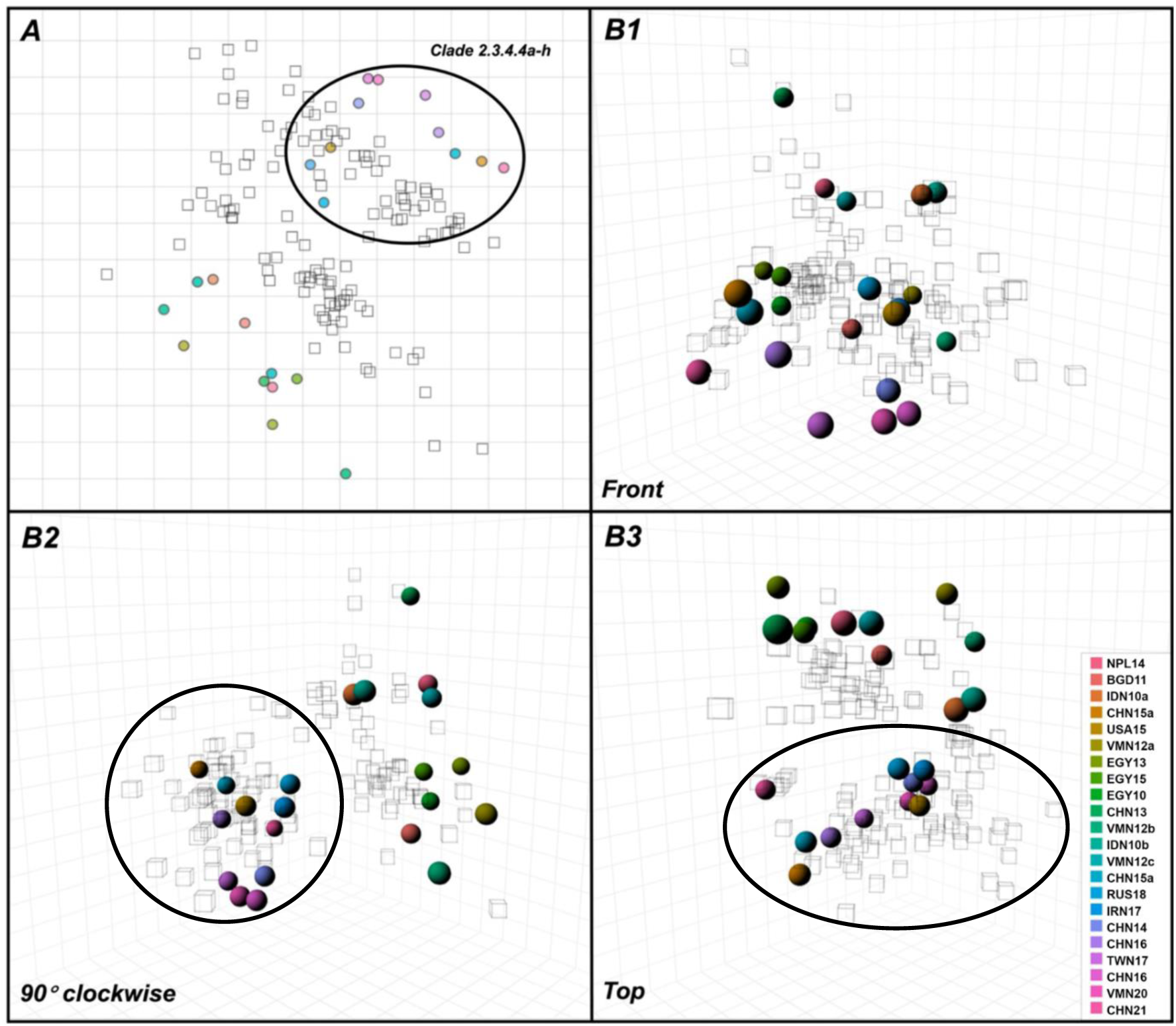
Antigenic cartography visualising serological relationships (GMTs) from **Table 3**. Maps plotted in 2- (**A**) and 3-dimensions (**B1-3**). Circles/spheres are antigens, coloured according to the key within **B3**; squares/cubes are sera. The grid plot represents one antigenic unit per square i.e., 2log_2_ HI titre. The 2.3.4.4 clade is circled to indicate it’s distinction. Maps were constructed in RStudio, using the Racmacs package.

The 2D and 3D antigenic maps clearly distinguished clade 2.3.4.4, which was visually segregated from the other clades. The antisera (represented as squares/cubes) from clade 2.3.4.4 largely clustered around the antigens (circles/spheres) of the same clade, whereas antisera from other clades were positioned more centrally on the map, situated between the 2.3.4.4 cluster and the remaining clade viruses. Additionally, the distribution of viruses within clade 2.3.4.4 revealed substantial intra-clade antigenic diversity.

### Comparative analysis of genetic and antigenic diversity highlighted putative antigenic residues driving variability between H5Nx clades

Antigenic distances derived from both 2D and 3D antigenic cartography maps were plotted against the corresponding amino acid variances in the highly variable HA1 region (Figure 5). Instances where large antigenic differences corresponded to small genetic differences suggest that a limited number of residue changes significantly impact antigenicity. To quantify this relationship, the ratio of antigenic distance to genetic distance was calculated for each antigen pair (Table S2). The pairs with the highest ratios (n = 28) were highlighted on the scatter plots, with shared pairs between 2D and 3D maps coloured in red, and pairs unique to the 2D or 3D maps coloured orange and purple, respectively.

**Figure 5:**
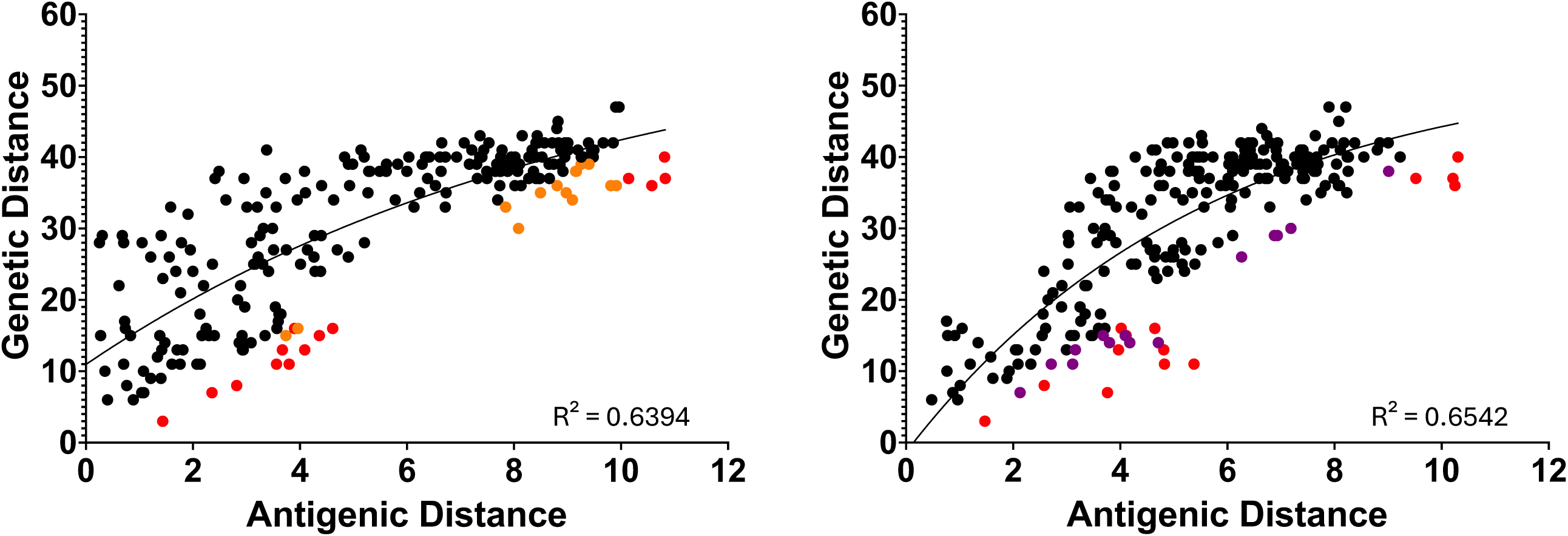
Scatterplots of HA1 genetic and antigenic distance pairwise comparisons between study strains. The comparisons with the greatest ratio (antigenic distance/genetic distance) (n = 28) are coloured on the plots. Antigenic distances from the 2-dimensional (2D) (**A**) and 3-dimensional (3D) (**B**) antigenic cartography maps (Figure 2). The same pairwise comparisons identified in both 2D, and 3D maps (n = 14) are highlighted in red and unique comparisons (n = 14 per) in orange and purple, respectively. Plots are fitted with non-linear, one- phase association lines of fit, with R² values indicated.

Both plots shared 14 out of 28 (50%) viral pairings, with the remaining 14 pairings unique to each plot. This difference in pair matching reflects the influence of the z-axis in the 3D plot on the overall distances between points. To determine which plot offered the most robust representation of the data, a non-linear, one-phase association curve was fitted to each, with the coefficient of determination (R²) reported. The 2D and 3D models yielded R² values of 0.6394 and 0.6542, respectively.

Residue variances in the HA1 region between the virus pairs were identified and highlighted (**Table 4**). Given its central position on the antigenic map, RUS18 was selected as the template HA for site-directed mutagenesis. Putative residues were mutated from the original amino acids in RUS18 to the alternatives predicted from paired comparisons. Additionally, the prevalence of each residue variant within the study strains was highlighted.

**Table 4:**
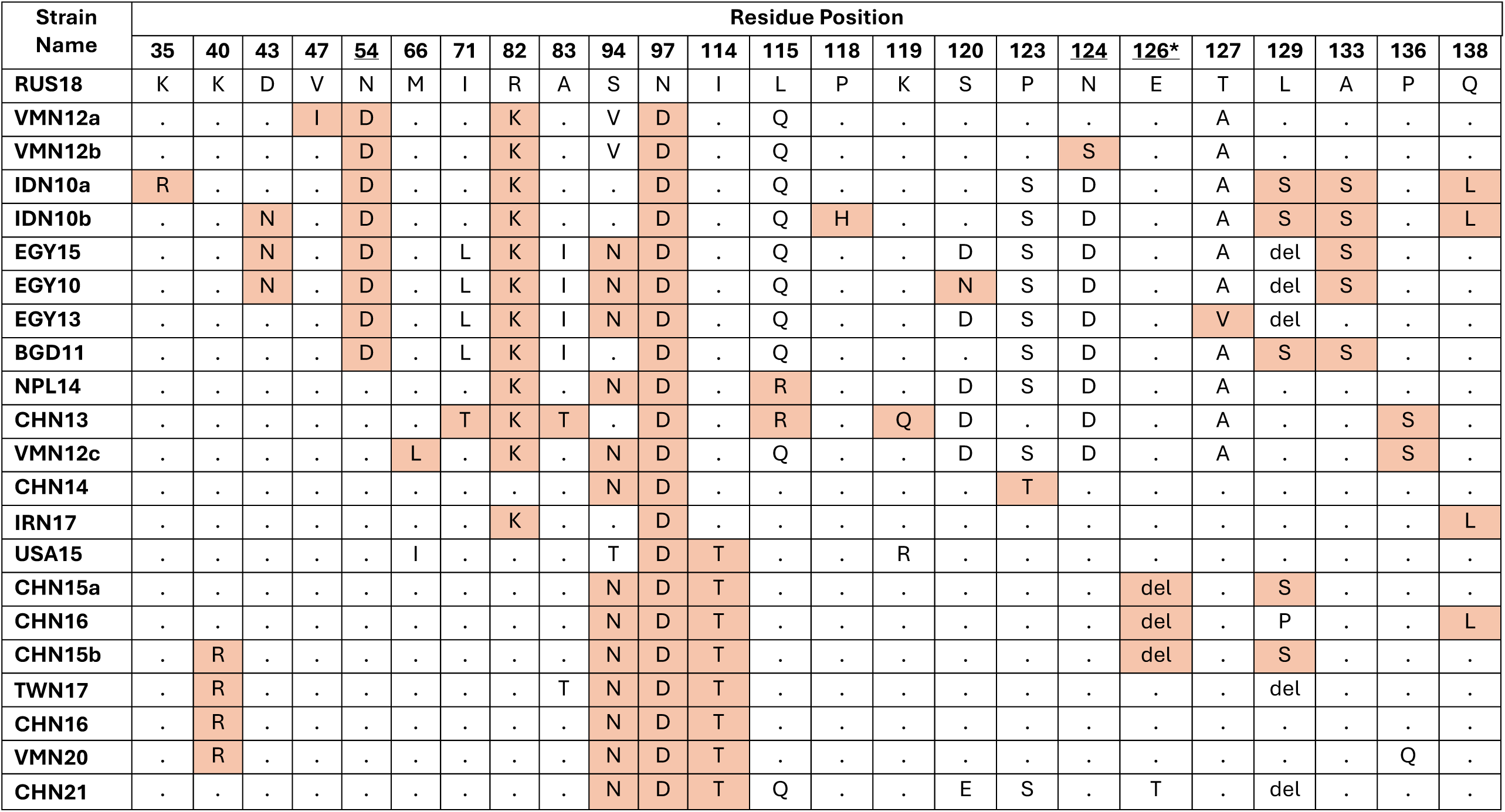
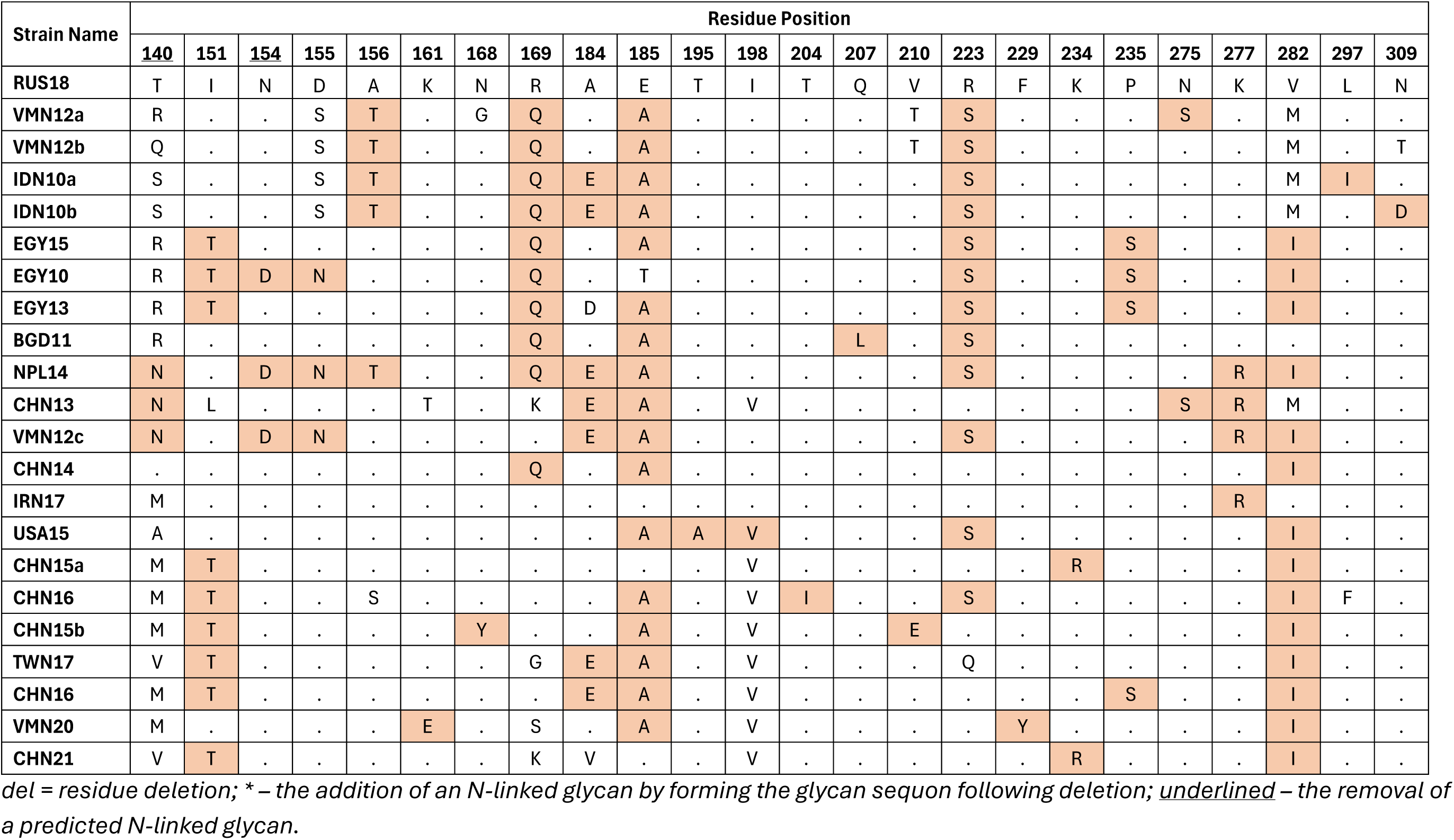
Putative antigenic residue selection and abundance in H5Nx study strains. Mutations highlighted in orange, with differences shown and identical residues ( . ) in reference to A/Chicken/Cheboksary/854/2018 (RUS18). Other variances within the putative position are unhighlighted.

From the 28 pairwise comparisons, 48 amino acid variations within the HA1 region were identified. Of these, 21 out of 48 (43.8%) variations were unique to a single strain, while the remaining variations were observed across multiple strains in this study **(Figure 6)**. Among these residues, four corresponded to predicted alterations in N-linked glycosylation sites, including one instance where an amino acid deletion introduced a new N-linked sequon.

**Figure 6:**
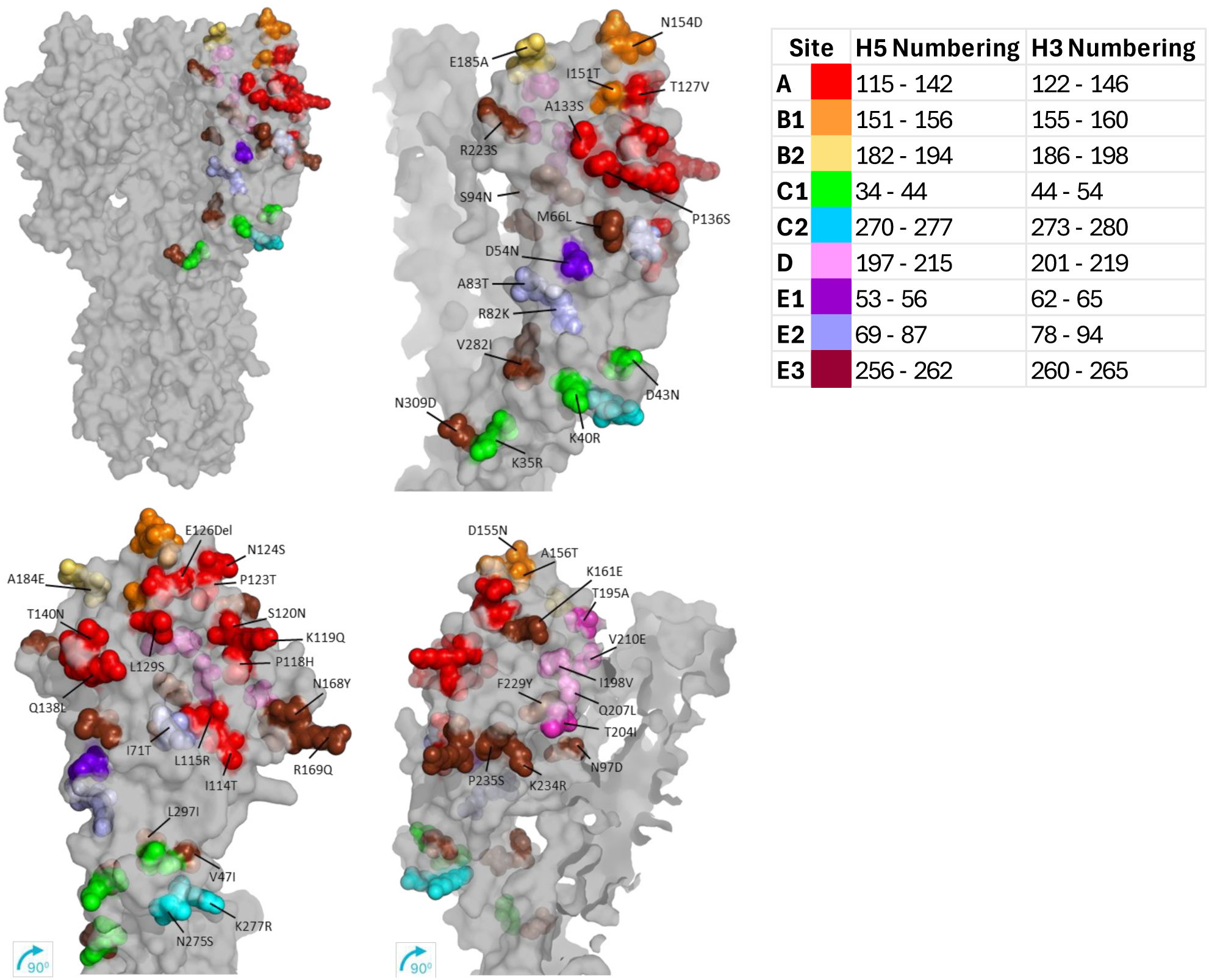
Protein model of the putative residue locations on A/Chicken/Cheboksary/854/2018 haemagglutinin (HA). Putative residues are displayed as spheres on the model surface, coloured by estimated antigenic site location [7, 9], or coloured brown if no corresponding site applies. Models are turned 90° clockwise and labelled for best visualisation. Models constructed and edited in Swiss-Model [3, 10] (using PDB: 6pcx.1 as template) and PyMOL [11]. Mature numbering used for site location, with H5 numbering for putative residues.

Subsequently, the putative residues were mapped onto a monomeric HA protein model to evaluate their location relative to known antigenic sites and overall structural position. Residues buried within the HA structure are less likely to result from immune escape and were therefore excluded from further consideration. Each putative antigenic epitope was then highlighted and colour-coded based on its predicted antigenic site using the H5 HA model (PDB ID: 6pcx.1) [23] **(Figure 6)**.

The antigenic sites of H5 are spread within the HA1 of the HA. Sites A, B and D are located within the receptor binding site (RBS), with A and B surrounding the receptor binding pocket. Sites E and C2 are distal to the RBS, within the less variable vestigial esterase domain at the base of the globular head. Site C1 is the most distal region, forming a concentrated region at the top of the stem and is the first site within the open-reading frame (ORF).

All putative epitopes were on the surface of the HA or protruded from the protein, including those outside of a previously determined antigenic region [23], and so were taken forward for further investigation.

### Antigenic characterisation of putative antigenic epitopes

Putative antigenic residues were introduced into the template HA, RUS18, via site-directed mutagenesis, with residues in close proximity mutated simultaneously. The 48 individual residue changes were consolidated into 29 mutagenesis reactions. Following confirmation by Sanger sequencing, the mutated HAs were rescued as recombinant viruses and evaluated for their impact on antigenicity using the HI assay with sera raised against the wild-type RUS18 HA. The changes in antigenic titres of the homologous antisera against each mutant virus were plotted as averages from three technical repeats and individual biological repeats (n = 7) **(Figure 7)**. A change in HI titre of ≥ 2log_2_ (equivalent to a 4 HAU difference) was used as the threshold to classify an antigenic variant.

**Figure 7:**
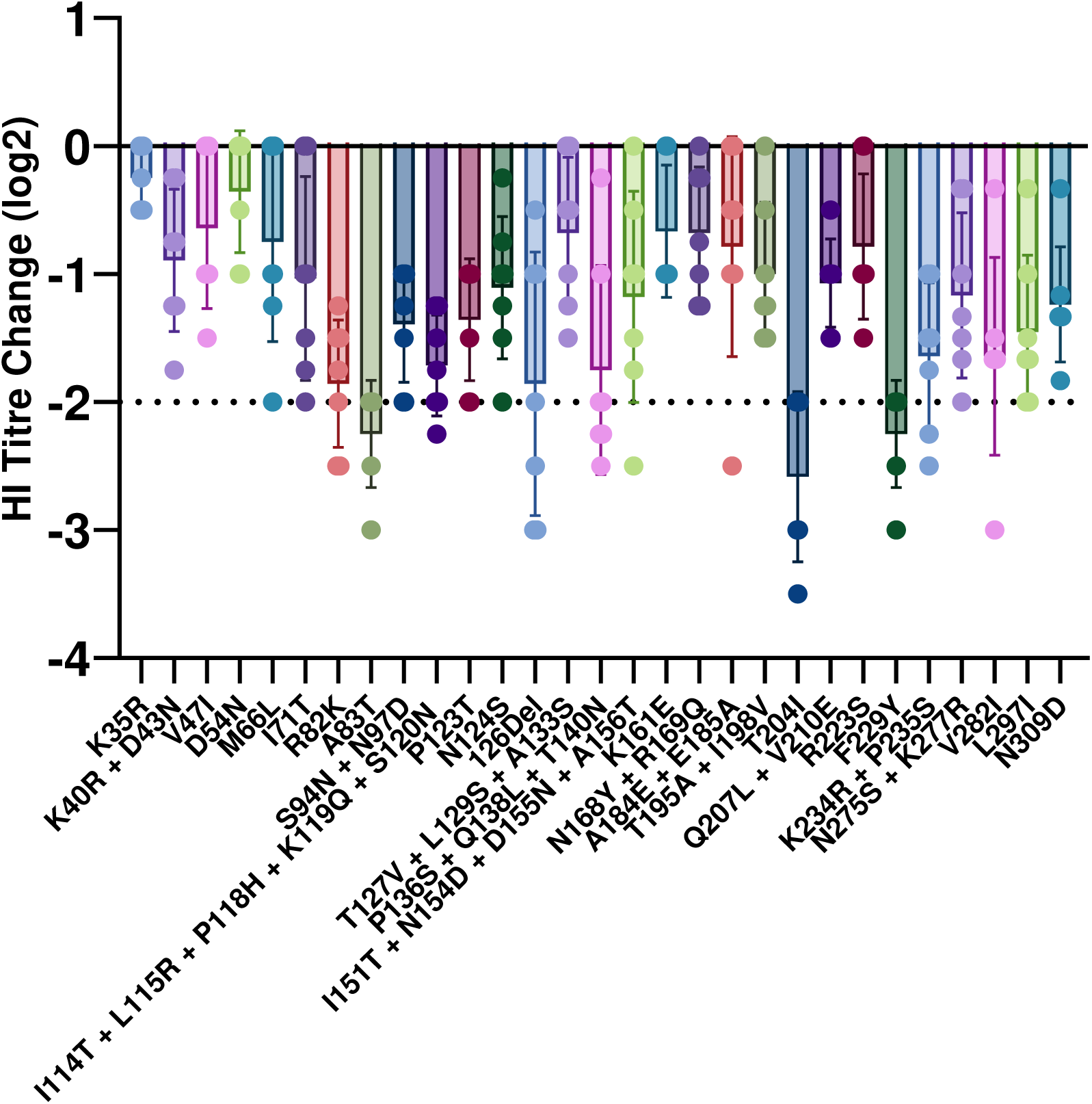
Geometric mean titre (GMT) change by haemagglutination inhibition (HI) of putative antigenic residues to previously homologous whole virus antisera. Titre changes greater than 2log_2_ are considered a significant antigenic change. The residues are indicated by the original amino acid, the position, and the amino acid change. Plots are individual bird sera and bars the range of standard deviation. Each plot GMT is the combination of three technical repeats.

Each putative residue mutation or combination influenced antigenicity to varying degrees, with three mutations A83T, T204I, and F229Y exceeding the antigenic variant threshold, showing GMT changes greater than 2log_2_. Among the mutants, 18 (62.1%) exhibited individual sera titre changes of 2log_2_ or more, although their GMTs did not surpass the threshold. Notably, mutations A83T, 126del, T204I, F229Y, and V282I induced sera titre changes equal to or greater than 3log_2_. The data also revealed substantial individual variation in neutralising antibody responses, with some mutant titres differing by up to 2.5log₂ between birds vaccinated with the same antigen, suggesting considerable heterogeneity in immune response despite identical antigen exposure and vaccination conditions.

## Discussion

The control of H5Nx AIVs in endemic regions relies heavily on vaccination strategies aimed at reducing poultry morbidity and mortality and limiting viral transmission. However, the recent unprecedented global spread of H5Nx AIVs, particularly those belonging to clade 2.3.4.4b has heightened the demand for vaccine interventions not only in endemic regions but also for the protection of poultry and vulnerable wild bird populations worldwide. Despite the dominance of clade 2.3.4.4b, co-circulation with other antigenically distinct lineages, including endemic clades 2.3.2.1a, 2.3.2.1c, and other subclades of 2.3.4.4, continues to pose a significant challenge [24, 25]. The success of vaccination campaigns is hindered by both the antigenic diversity among circulating strains and mismatches between vaccine seed strains and field viruses. Therefore, this study aimed to identify specific amino acid residues that contribute to antigenic differences between recently or currently circulating H5Nx clades, which may compromise vaccine efficacy and inform vaccine seed strain selection. While genetic evolution is typically characterised through phylogenetic analysis [26], antigenic evolution can be more directly assessed using antigenic cartography derived from HI assay data [27]. Accordingly, both phylogenetic and antigenic mapping approaches were employed in tandem to identify and evaluate putative residues driving antigenic diversity.

Clade-representative viruses were selected, and their classification confirmed through phylogenetic analysis, which also highlighted key monophyletic amino acid signatures characteristic of each clade and subclade. Notably, clade 2.3.4.4 was greatly distinguished with substantial inter- and intra-clade genetic variability, with up to 39 amino acid intra-clade differences observed. de Vries *et al.* [28] previously identified seven characteristic mutations within the HA protein of clade 2.3.4.4 (K82R, T156A, N183D, K218Q, S223R, N240H, and A263T). All but one (S223R) were consistently conserved across 2.3.4.4 subclades; S223R appeared to have been lost in subclades 2.3.4.4c to 2.3.4.4h. Interestingly, A263T, although conserved within clade 2.3.4.4, also appeared across several other clades (1 to 2.2.2.1), with notable exceptions in clades 2.1.3.2, 2.3.2.1a – c, and 2.4. Across the dataset, residue 263 exhibited a roughly 50:50 distribution of alanine and threonine. Furthermore, 12 additional residues were conserved within HA1 across all 2.3.4.4 viruses, with two more residues conserved specifically in subclades 2.3.4.4c – h. Collectively, clade 2.3.4.4 exhibited the most conserved antigenic markers, followed by clade 2.3.2.1 another genetically and antigenically distinct clade frequently implicated in recent poultry outbreaks [10, 29]. These findings highlight the profound antigenic and genetic divergence of clade 2.3.4.4 and the resulting complexity in formulating a broadly protective H5 vaccine.

Another immune escape mechanism employed by influenza A viruses involves the acquisition of N-linked glycosylation sites near antigenic regions. The addition of bulky oligosaccharide moieties can sterically hinder antibody access to epitopes. Apart from the glycosylation site at residue 165, five conserved glycosylation sites were identified across all strains, located within the stem domain of HA. These stem glycans are critical for maintaining the structural integrity and fusion functionality of HA [30]. In contrast, glycosylation sites within the globular head are highly variable and contribute to immune evasion [31] and modulation of receptor binding [32]. Frequently occurring glycosylation sites at positions 10, 11, 23, 154, 165, 193, 286, and 484 have been previously reported across multiple H5 clades and are thought to stabilise the protein or shield antigenic epitopes [33, 34]. Less common glycosylation sites at positions 70, 140, 182, and 289 were observed sporadically, likely reflecting strain-specific adaptations. These variable glycans, often located within or near antigenic sites in the globular head region of HA, may arise in response to immune pressure or serve to enhance receptor binding affinity.

Importantly, genetic diversity does not always correlate with antigenic diversity. Even minor amino acid substitutions in HA can lead to substantial antigenic shifts. To evaluate the antigenic cross-reactivity of selected H5 strains, each HA gene was rescued into a low pathogenicity virus backbone using PR8 internal and NA genes, and polyclonal antisera were raised in chickens. Antigenic relationships were assessed HI assays. Homologous antigen–sera pairs produced titres well above protective thresholds, confirming successful inoculation and seroconversion. While there was a general correlation between genetic and antigenic variation, clade 2.3.4.4 again demonstrated marked antigenic heterogeneity. Consistent with previous findings, clade 2.3.4.4 viruses were poorly neutralised by monoclonal or polyclonal antibodies raised against other clades, including those co-circulating within the same geographical regions [35]. The rapid and ongoing diversification of 2.3.4.4 subclades has been widely documented, most notably, the persistence of clade 2.3.4.4b in wild bird populations across Europe, the Americas, and Africa, and the endemic circulation of clades 2.3.4.4d-h in Asian poultry [36]. This independent evolution within each subclade is driving further antigenic divergence. Consequently, the subclades are becoming antigenically distinct entities, potentially warranting a reassessment of the current H5 clade nomenclature to better align with phylogenetic and antigenic boundaries [37].

Antigenic cartography enables the visualisation of antigenic evolution and the relationships between viral strains and corresponding antisera in two- and three-dimensional space. HI titres, generated by assessing the cross-reactivity of heterologous antisera to recombinant viruses, were plotted relative to their similarity to homologous sera titres. A key observation from both maps was the clear segregation of clade 2.3.4.4a–h antigens from all other clades (1.1 to 2.3.2.1b). In line with the genetic analysis, clade 2.3.4.4 exhibited marked antigenic uniqueness, demonstrating minimal cross-reactivity with other clades and confirming its pronounced antigenic divergence. Furthermore, the dispersion of antigen plots within clade 2.3.4.4 revealed considerable intra-clade antigenic variation. These findings align with multiple recent studies reporting the pronounced antigenic distinctiveness of clade 2.3.4.4, including the currently dominant subclade 2.3.4.4b, which exhibits notable divergence even within its own lineage [36, 38, 39].

The virus pairs exhibiting the highest antigenic-to-genetic distance ratios, indicating substantial antigenic divergence with minimal amino acid variation, were selected for further analysis. Amino acid differences between these virus pairs were identified, yielding a list of putative residues for downstream analysis. Notably, the highest antigenic/genetic ratios were observed between viruses from closely related clades (e.g., 2.2.1 and 2.2.1.2, or 2.3.4.4d and 2.3.4.4e). Several amino acid substitutions were recurrent across multiple strains, particularly among those belonging to similar clades or subject to shared geographical constraints. As anticipated, the majority of variations were concentrated in the globular head domain of HA1 (amino acids 42–272), a region commonly implicated in antibody recognition. To differentiate putative antigenic epitopes from synonymous or structurally buried mutations, each candidate residue was mapped onto the HA monomer model of RUS18. Mutations not exposed on the surface of the protein were excluded, as their likelihood of contributing to antibody escape is low. Surface-exposed residues were further analysed in the context of defined antigenic regions. For H5 HAs, five principal antigenic sites (A – E) have been described based on their spatial distribution across the HA head. While consideration of these canonical regions helps reduce false positives [40], this study also aimed to identify novel epitopes located outside these predefined sites. This approach is supported by reports of escape mutations arising beyond the established antigenic regions, often near the receptor-binding site or adjacent to residues known to critically influence receptor affinity factors that may confound HI assay interpretations [41, 42].

A total of 48 putative antigenic residues were identified, of which 34 have been previously reported as antigenic epitopes [41–53] while 16 were either novel or located outside established antigenic sites. Of the 32 residues situated within defined antigenic regions, their distribution was as follows: Site A (n = 14), Site B (n = 7), Site C (n = 3), Site D (n = 3), and Site E (n = 5). The most influential epitopes were predominantly located in or near the highly variable and well-characterised receptor-binding domain (RBD) of HA. Site A includes the critical 130-loop (H5 numbering: residues 131–134), while Site B encompasses both the 150-loop (residues 151–159) and the 190-helix (residues 186–194), all structural motifs essential for receptor affinity, specificity, and host range [54, 55]. Site D contributes the final RBD element, the 220-loop (residues 217–224), which also influences receptor specificity, although to a lesser extent than Sites A and B.

In contrast, Sites C and E do not overlap with the structural components of the RBD but instead coincide with the vestigial esterase and fusion domains of HA. Antibodies targeting the RBD often exert potent neutralising activity and tend to be strain- or clade-specific, contributing to rapid immune-driven selection at these sites. Conversely, antibodies directed against the vestigial esterase domain are typically non-neutralising but may confer cross-clade protection by engaging Fc-mediated effector functions such as antibody-dependent cellular phagocytosis (ADCP) and antibody-dependent cellular cytotoxicity (ADCC) [56–60]. Each putative residue was introduced into a candidate HA (RUS18) for investigation of antigenic influence using previously homologous antisera. RUS18 was chosen due to its central positioning within the antigenic cartography maps, low saturation of N-linked glycans as some were to be removed and introduced, which may distort intracellular trafficking, and relevance to currently dominating strains. Of the 29 mutants, 18 (62.1%) achieved individual values over or equal to a 2log_2_ titre change but did not surpass the threshold by GMT. However, A83T, 126del, T204I, F229Y and V282I, each induced sera titre above or equal to a 3log_2_ change. These mutations were within 2.3.2.1b, 2.3.4.4d, 2.3.4.4d, 2.3.4.4g, and both 2.2.1 and 2.3.4.4a-h except for 2.3.4.4b, respectively. A83T, 126del and T204I mutations are situated within antigenic sites E, A and D, respectively, with F229T and V282 situated outside of an assigned site.

Three residues A83T, 126del, and V282I are recognised antigenic epitopes. Residue 83 was identified through peptide mutagenesis studies, in which its substitution resulted in the loss of neutralisation by monoclonal antibodies, confirming its antigenic relevance [46]. This study supports and complements those earlier findings. The 126 site has been reported to acquire N-linked glycosylation, effectively masking an epitope targeted by monoclonal antibodies. This modification not only reduces virulence in mice potentially due to increased susceptibility to serum inhibitors but has also emerged as an antigenic escape mutation [50, 61]. In agreement with Kaverin *et al*. [61], the loss of glycosylation at this site in the present study was associated with reduced cross-reactivity to antisera and significantly decreased HI titres. Similarly, V282I has been previously implicated in antigenic drift and identified as a monoclonal antibody escape mutation [43].

In contrast, T204I and F229Y have not been directly characterised as epitope-defining mutations. However, residue 204 resides within the highly antigenic site D, proximal to the receptor-binding domain (RBD), and is generally conserved as threonine across clades. Although residue 229 does not fall within any defined antigenic site, it is structurally adjacent to residue 204 (**Figure 6**), suggesting it may exert an indirect influence on local antigenicity through conformational effects or epitope shielding.

### Conclusion

The dominant and currently circulating clade 2.3.4.4 is both genetically and antigenically distinct from predecessor clades, exhibiting substantial intra clade variability. Through detailed mapping of the highly variable HA1 region, a combination of antigenic and non-antigenic residues was identified that differentiate clades. Of the mutations assessed, 18 (62.1%) induced a ≥2 log₂ change in HI titre individually but did not exceed the threshold when evaluated by geometric mean titre (GMT). In contrast, five mutations A83T, 126del, T204I, F229Y, and V282I, each resulted in ≥3 log₂ reductions in HI titre, highlighting their potential as key antigenic determinants. Notably, substantial variability in titres was observed between individual birds immunised with the same antigen, with some mutants displaying up to 2.5 log₂ differences, underscoring the impact of host-specific factors on the neutralising antibody repertoire.

This integrative approach, combining genetic and antigenic data, enhances our understanding of the antigenic evolution of H5 HA, particularly within subclade 2.3.4.4b. The identification of residues that drive or fail to drive antigenic drift provides valuable insight for refining vaccine design. These findings support the development of more effective immunogens for poultry and in light of recent zoonotic spillovers, potentially for use in mammals, including humans.

## Materials and Methods

### Acquisition and modification of sequences

Virus strains were selected as representatives of recently or currently active clades of H5Nx Gs/Gd lineages (clade 1 – 2.3.4.4h). Representative strains from clades 1.1.1 to 2.3.2.1c were selected based on the 2014 updated nomenclature published by the WHO/OIE/FAO [14] while strains from clades 2.3.4.4a and 2.3.4.4d to 2.3.4.4h were selected according to a study by Bui *et al* [15] for the Centers for Disease Control and Prevention (CDC). Complete HA sequences of the selected strains were downloaded from the NCBI (National Center for Biotechnology Information) [16] and GISAID (Global Initiative on Sharing All Influenza Data) [17] databases.

Downloaded HA sequences were inspected for nucleotide errors or frameshifts. Sequences were visualised and aligned using MUSCLE [62], and clustering was performed using UPGMA [63] in MEGA (version 10) [64]. Nucleotide sequences were trimmed to the open reading frame (ORF) and translated to ensure correct structural formation. In cases of frameshifting or sequence errors, alignments were reviewed, and the most common nucleotide at the affected position was introduced to correct the sequence.

Untranslated regions (UTRs) complementary to the pHW2000 expression system were added to both the 5′ and 3′ ends of the sequences. These UTRs included gene- and subtype-specific segments along with BsmBI (Esp3I) restriction enzyme sites [65, 66]. The translated region was further modified by replacing the polybasic amino acid cleavage site with a monobasic site to remove high virulence, enabling experimentation at containment level 2. The entire sequence was also examined for additional BsmBI recognition sites, which were removed through synonymous codon substitutions. The modified HA sequences were synthesised by GeneArt Gene Synthesis (Thermo Fisher Scientific) and cloned into the pMT vector by the manufacturer.

### Gene subcloning

The pMT vectors containing HA gene inserts and empty pHW2000 vectors were digested at the designated restriction sites within the UTRs using the BsmBI enzyme and CutSmart® Buffer (NEB), according to the manufacturer’s instructions. The pHW2000 vector was additionally dephosphorylated prior to ligation using Antarctic Phosphatase (NEB). HA inserts were isolated by gel electrophoresis and extracted using the QIAquick Gel Extraction Kit (Qiagen). The digested HA genes and digested/dephosphorylated pHW2000 vector were then ligated using the T4 DNA Ligase Kit (NEB), following the manufacturer’s instructions.

### Bacterial transformation and plasmid expansion

Ligation products were transformed into in-house-prepared chemically competent *Escherichia coli* (DH5-α, NEB) using the standard heat-shock protocol [67]. For plasmid sequencing by Sanger, plasmids were extracted using the QIAprep Spin Miniprep Kit (Qiagen), following the manufacturer’s instructions. For preparation of working stocks, plasmids were extracted from larger bacterial cultures using the QIAprep Spin Maxiprep Kit (Qiagen), also according to the manufacturer’s instructions.

### Generation of recombinant influenza viruses

The well-established 8-plasmid bidirectional reverse genetics (RG) system was used to generate recombinant H5Nx viruses, incorporating the HA and NA genes from H5Nx strains of interest and the internal genes from the lab-adapted H1N1 strain A/Puerto Rico/8/1934 (PR8) [68–71]. Virus stocks were expanded by *in ovo* propagation using 10-day-old embryonated White Leghorn chicken eggs (VALO). Eggs were candled daily to monitor embryo viability. Propagation was terminated by chilling at 4 °C for a minimum of four hours. Following successful viral propagation, viral RNA (vRNA) was extracted to confirm the HA and NA sequences. Extraction was performed from allantoic fluid using the QIAamp Viral RNA Kit (Qiagen) and the centrifugation protocol, following the manufacturer’s instructions.

### Haemagglutination assay and haemagglutination inhibition (HI) assay

Standard influenza virus haemagglutination assays were performed to titrate virus stocks and assess viral activity based on haemagglutinin binding. These assays are applicable to both active and inactivated influenza viruses [72, 73]. Briefly, haemagglutination inhibition (HI) assays were carried out using four haemagglutinating units (HAU) of each virus, incubated with 2-fold serial dilutions of antiserum raised in specific-pathogen free (SPF) White Leghorn chickens. HI titres were expressed as the reciprocal of the highest antiserum dilution at which haemagglutination was completely inhibited.

### Raising antisera

Stocks of recombinant viruses were prepared in embryonated chicken eggs and subsequently inactivated using 0.1% (v/v) β-propiolactone (BPL) (Sigma-Aldrich). To confirm successful inactivation, 200 µl of each virus preparation in allantoic fluid was inoculated into embryonated eggs across three consecutive passages, with each subsequent egg inoculated using allantoic fluid from the previous passage. Virus replication was assessed after each passage by haemagglutination assay and recorded as haemagglutination units (HAUs). Any virus preparations testing positive for haemagglutination activity required repeated inactivation and passaging.

To achieve the desired viral titres of 1024 HAU/dose of vaccine, the allantoic fluid was concentrated via ultracentrifugation. Vaccines were formulated at a 30:70 antigen-to- adjuvant ratio using MONTANIDE™ ISA 71 VG adjuvant (SEPPIC), following the manufacturer’s instructions.

Day-old White Leghorn chickens were inoculated subcutaneously at the back of the neck with a single 200 µl vaccine dose and similarly boosted at 21 days post-vaccination. Birds were bled according to their respective weights at 7, 14, 21-, 28-, 35-, and 42-days post-vaccination, with final bleed at day 42 marking the end of the experiment.

### TCID_50_ and microneutralisation assay

The amount of live virus in the allantoic fluid was estimated using TCID₅ ₀ (Tissue Culture Infectious Dose 50%) assays for subsequent use in microneutralisation (MN) assays. Confluent monolayers of Madin-Darby Canine Kidney (MDCK) cells, seeded into 96-well plates, were washed with PBS and infected with 2-fold serially diluted virus preparations in quadruplicate. After a 1-hour incubation at 37°C, the virus inoculum was removed and replaced with DMEM supplemented with TPCK-trypsin (Sigma-Aldrich). Plates were incubated for 72 hours, after which the cells were stained with crystal violet solution. Viral titres were calculated using the Spearman–Karber method [74].

MN assays were performed using 150 TCID₅ ₀ of each virus per well. MDCK cells were seeded as described above. Chicken antisera were heat-inactivated at 56°C for 30 minutes, then 2-fold serially diluted in DMEM. The diluted antisera were incubated with virus in triplicate wells of a separate 96-well plate for one hour at 37°C. Prepared MDCK cells were washed with PBS, and the virus-antisera mixtures were added to the wells and incubated for a further 72 hours at 37°C. Cells were then fixed and stained with crystal violet solution to assess the neutralising activity of anti-influenza antibodies present in the antisera.

### Site-directed mutagenesis

Putative antigenic epitopes were introduced into the H5 HA of A/chicken/Cheboksary/854/201 (used a backbone template) by site mutagenesis, using the QuikChange Lightning Site-directed Mutagenesis kit (Agilent), following the manufacturer’s instructions. 18 overlapping primer pairs were designed using the Agilent Primer Design (Agilent) for optimal primer design **(Table S3)**.

### Phylogenetic analysis

Phylogenetic analysis of nucleotide sequences was performed using IQTree [75, 76]. Trees were initially constructed as Neighbour-joining to predict the most accurate tree model for Maximum-likelihood [77]. The classification and assignment of clades of H5Nx viruses assumes a common ancestral node and monophyletic evolution with a bootstrap value ≥ 60 at the defining node following 1000 bootstrap replicates [14]. Trees were annotated and coloured using FigTree [78].

### Antigenic cartography

Antigenic cartography was used to visualise antigenic relationships and cross-reactivity among a panel of influenza virus antigens and corresponding antisera, based on haemagglutination inhibition (HI) assay titres [27]. The maps employ the theoretical framework of “shape space” [79] in which antigenic distances reflect differences in antibody recognition, with shorter distances indicating higher antigenic similarity often due to structural similarity or shared epitopes. Antigenic maps were constructed and analysed using Acmacs [22] and RStudio, incorporating the Racmacs package and associated tools. A modified multidimensional scaling (MDS) algorithm was applied to transform HI titres into Euclidean distances, minimising error to best fit the observed serological data.

### Development of putative antigenic epitopes

To identify key antigenic epitopes that may drive changes in virus antigenicity and impact vaccine efficacy, antigenic distances derived from antigenic cartography were compared pairwise with the corresponding genetic distances, measured as amino acid residue differences, following the approaches described in [27, 80]. Virus pairs exhibiting the greatest antigenic distance but the smallest genetic distance, thus yielding the highest antigenic/genetic ratio were prioritised for further analysis. Amino acid differences between these virus pairs were considered as putative antigenic determinants and were subsequently introduced into a template HA sequence via site-directed mutagenesis for functional investigation.

### Lines of regression

Non-linear regression models were fitted to the pairwise comparisons of genetic and antigenic distances. The model with the highest R-squared value, indicating the greatest proportion of explained variance between the variables, was selected.

### Ethics

All animal studies and procedures were conducted in accordance with European and United Kingdom Home Office regulations, including the Animals (Scientific Procedures) Act 1986 Amendment Regulations 2012, under approved project licences (P68D44CF4 and PP6471846) issued by the UK Home Office. All experimental procedures were reviewed and approved by the Animal Welfare and Ethical Review Board at The Pirbright Institute and were performed by highly trained and licensed personnel. Animal welfare was closely monitored throughout the studies, with adherence to the 3Rs principles wherever possible.

## Acknowledgments

The work described herein was funded by UK Research and Innovation (UKRI) Biotechnology and Biological Sciences Research Council (BBSRC) grants ISPG, T BBS/E/PI/230001C, BBS/E/PI/230001A, BBS/E/PI/23NB0003, BB/Y007298/1, BB/W003325/1, BB/T013087/1, and the Global Challenges Research Fund (GCRF) One Health Poultry Hub (BB/S011269/1). The funders had no role in study design, data collection, data interpretation, or the decision to submit the work for publication. We also acknowledge the support of the animal services staff at the Pirbright Institute, whose help was instrumental in facilitating this research.

---

**Table S1:**
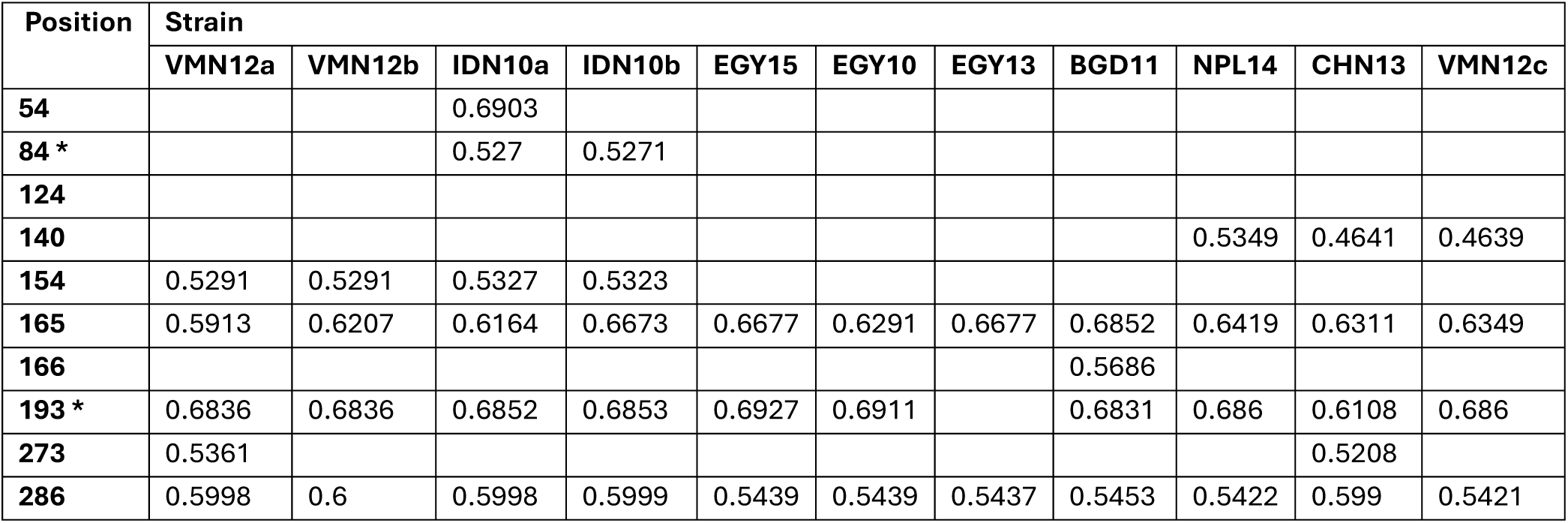
N-linked glycosylation prediction of H5 AIV HA sequences selected for this study. The ’potential’ score is the mean output of nine neural networks. Values above 0.5 with the sequon Asn-X-Ser/Thr without proline at X are deemed positive predictions. Values labelled with (*) possess proline at X and are negative.

**Table S2:**
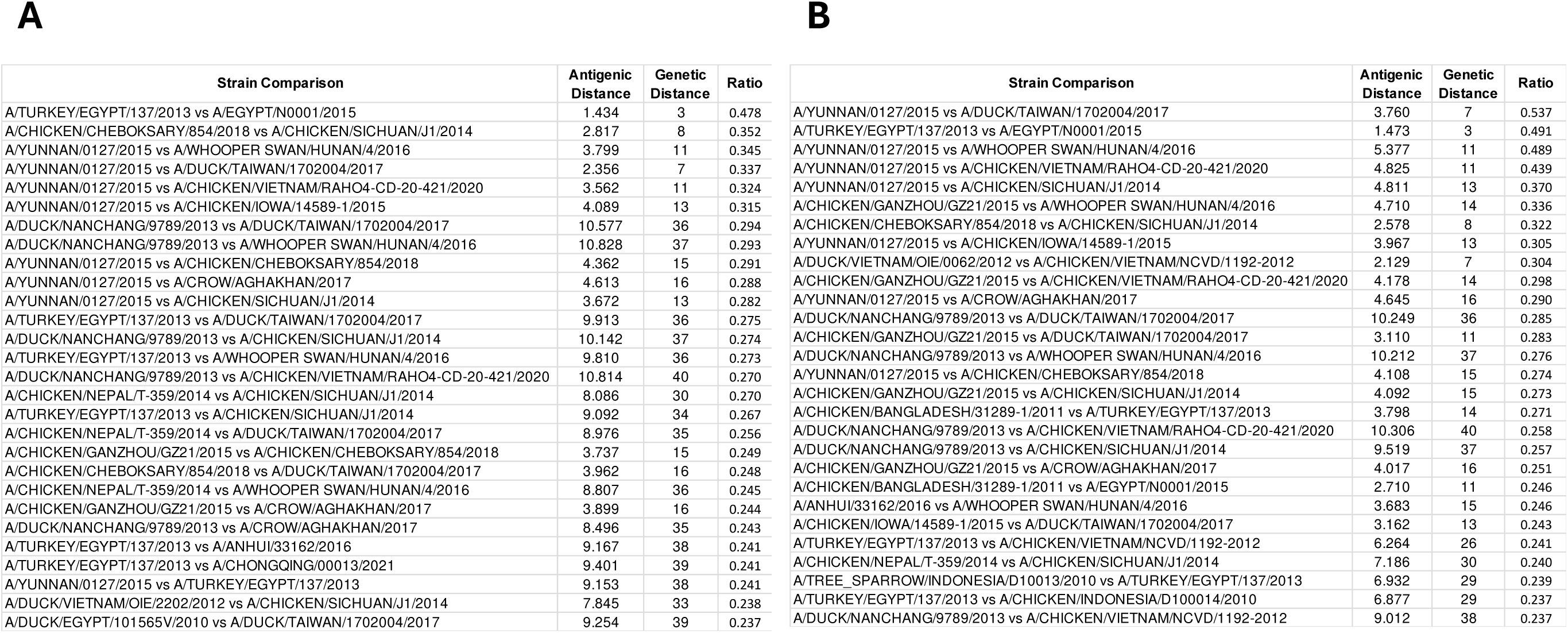
Pairwise comparisons of antigenic and genetic variances between the study strains, sorted in decreasing values of antigenic distance per genetic distance (ratio). Antigenic distances are the total antigenic units from 2-dimensional (**A**) and 3-dimensional (**B**) antigenic cartography. Genetic distance was calculated as the total amino acid variance between each HA1. Highest n=28 ratios for each dimension shown.

**Table S3:**
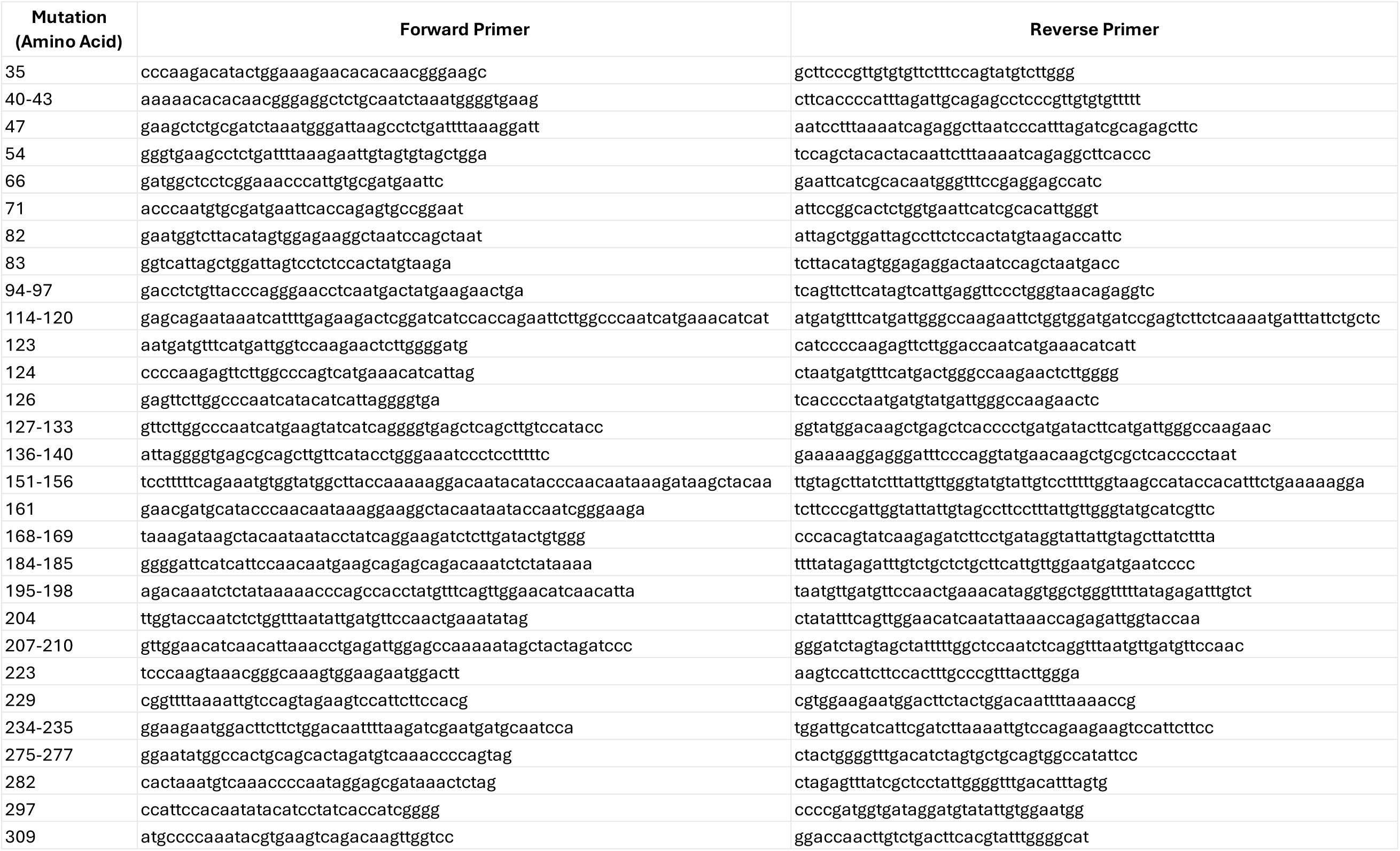
QuikChange Lightning Site-directed Mutagenesis (Agilent) Primer Pairs.

**Figure S1:**
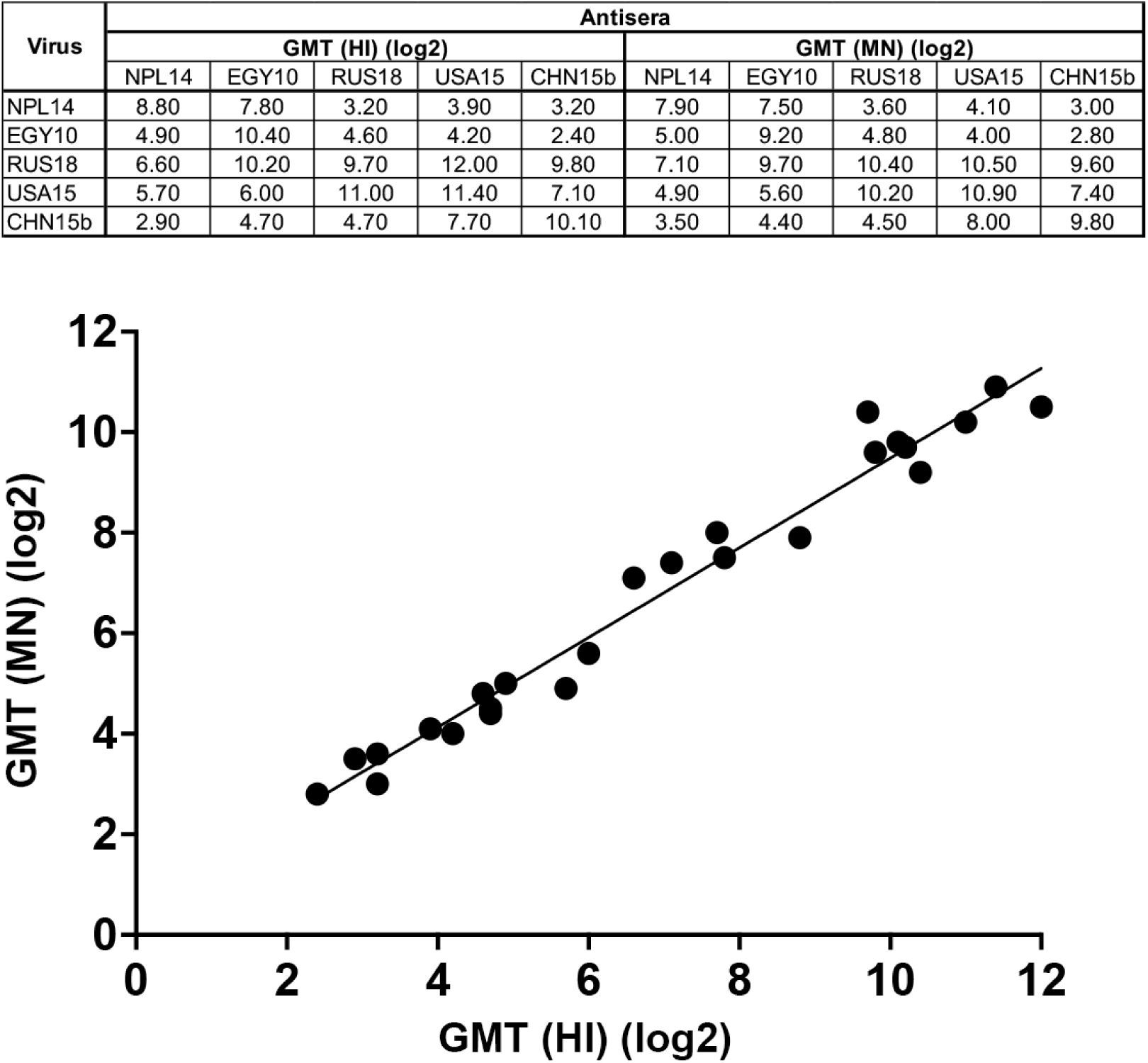
Comparison of the geometric mean titre (GMT) of haemagglutination inhibition (HI) and mircroneutralisation (MN) tests between NPL14, EGY10, RUS18, USA15 and CHN15b antigens and antisera, with viruses paired against homologous antisera in bold. Scatterplot of HI against MN titres, with a simple linear regression line with an R2 value of 0.9712.

